# Metrics for estimating individual-level changes in functional brain connectivity and their correspondence with topology changes

**DOI:** 10.1101/2025.06.12.639125

**Authors:** Katherine L. Bottenhorn, Jordan D. Corbett, Hedyeh Ahmadi, Megan M. Herting

## Abstract

With the rise of precision medicine and large neuroimaging datasets, measuring brain changes on an individual level becomes more possible and more important. However, functional connectivity has some mathematical and conceptual quirks that make estimating individual-level changes more complicated than, for example, estimating structural brain changes. Here, we compared six different change scores, i.e., metrics for quantifying change in functional connectivity, using a large sample with two time points, roughly 2 years apart, of low-motion data from the ABCD Study (N=2,719, ages 9-13 years). First, we compared distributions and potential interpretations of each metric. Then, we assessed how well different metrics captured topology change estimates by comparing them to individual-level changes in clustering coefficient, betweenness centrality, and network strength. As multivariate metrics may be more reliable than single connections and researchers often interpret connectivity changes in topological terms, this offers additional insight into the implications of change score choice. Overall, our analyses revealed widely different distributions between change scores that conferred vastly different results and interpretations of functional connectivity changes between time points. Thus, we provide recommendations for each change score and its optimal (or suboptimal) use cases, depending on the population, study design, and research question.

## Introduction

Functional magnetic resonance imaging (fMRI) provides a valuable, noninvasive window into *in vivo* brain function. Correspondence between the fMRI blood-oxygen-level-dependent (BOLD) signal of spatially disparate brain regions has been used to estimate the human brain’s functional connectivity (FC), revealing a functional architecture comprising several canonical, large-scale networks (Greicius et al., 2003; Smith et al., 2009) that persist across individuals (Gordon et al., 2017), states of consciousness (Boly et al., 2008), and age (Betzel et al., 2014; Vij et al., 2018). Within these canonical networks, however, subtle changes in functional topology have been observed during brain development (Fair et al., 2007; Gracia-Tabuenca et al., 2023; Grayson & Fair, 2017), aging (Betzel et al., 2014; Chan et al., 2014), disease, injury, treatment, and, more recently, even daily and weekly changes in behavior, hormones (Bottenhorn et al., 2024; Pritschet et al., 2020), and lifestyle factors (Poldrack et al., 2015). Thus, FC changes are broadly relevant phenomena in both health and disease, for both development and aging, to study treatment, training, injury, or lifestyle effects on brain function. As human neurosciences increasingly address differences in the brain, however, it becomes increasingly important to establish robust methods for assessing FC change on an individual level.

Identifying robust individual differences and changes in (FC) has been historically precluded by the preponderance of small, insufficiently powered human neuroimaging studies (Button et al., 2013; Marek et al., 2022; Yarkoni, 2009). However, the increasing number and size of neuroimaging datasets facilitates robust individual differences analyses and the advancement of individual-centric analytic strategies (Laird, 2021). Moreover, as recent work has emphasized the influence of individual-specific factors in large-scale brain networks (Gratton et al., 2018) and studies continue to highlight individual differences’ impact on brain-wide association studies (Bandettini et al., 2022; Marek et al., 2022), individual-centric analytic approaches have become more common and important for human neuroimaging research. Ideally, longitudinal designs with three or more data collection time points per participant allow for estimating individual-level slopes (e.g., from a linear mixed effects or general additive mixture model). However, repeated measures fMRI studies are often limited to two time points for either feasibility and/or by design to studying treatment, injury, and/or training effects on FC. However, FC has mathematical and conceptual properties that complicate assessments of its change across two time points. While there is substantial literature considering the merits of different metrics for computing FC, product-moment correlations of BOLD signals, and Z-scores thereof, remain the most commonly used metrics by which FC is estimated (Hlinkaa et al., 2011). Product-moment correlations range from *r* = −1, indicating perfect anticorrelation, to *r* = 1, indicating perfect correlation. If interpretations of FC were to reflect this mathematically, two brain regions whose BOLD signals exhibit a correlation of *r* = −1 might be construed as perfectly “disconnected” whereas *r* = 1 would reflect perfectly “connected” brain regions (Chai et al., 2012). Yet, when considering changes in correlation-based FC estimates, two pieces of information become important to consider: the *magnitude* of connectivity between regions (i.e., |*r*|) and the *nature*, or sign, of that connectivity (i.e., cooperative activations when *r* > 0 vs. opposing activations when *r* < 0). For example, a change from *r* = −0.5 to *r* = 0 calculated via simple subtraction is mathematically identical to a correlation from *r* = 0 to *r* = 0.5 or from *r* = −0.25 to *r* = 0.25, although in the context of FC these likely reflect very different functional changes with plausible differences in neural underpinnings and implications. Thus, the absolute value of *r* and its sign each contain valuable information about FC and about the nature of communication between brain regions, which can be easily lost when calculating changes.

To date, two studies have each leveraged two time points of resting state fMRI data to estimate FC changes in the ongoing, longitudinal Adolescent Brain Cognitive Development Study (ABCD Study®), a nationwide study that enrolled over 11,000 children ages 9-10 years in 2016 (Volkow et al., 2018). With these data, Saha et al. identified patterns of changing FC between ages 9 and 13 years that are related to psychiatric and cognitive factors via independent component analysis of the simple change in FC (i.e., *FC*_*T2*_ *-FC*_*T1*_; where *T*_*1*_ indicates the first data collection time point and *T*_*2*_ indicates the second) (Saha et al., 2024). Alternatively, Bottenhorn et al., described changes in large-scale network FC from ages 9 to 13 years old, and heterogeneity therein due to sex and puberty, estimated via annualized percent change (Bottenhorn et al., 2023) – a metric that has been used elsewhere in the neurodevelopmental literature (Mills et al., 2021). Although both studies assess FC changes across the same age range in the same sample, it is difficult to compare or contextualize their findings without knowing the effects of their different approaches to calculating FC change (i.e., simple change versus annualized percent change). To address this, the present study has profiled a number of different metrics for assessing changes in functional connectivity between two time points using tabulated resting state fMRI data from over 2,700 individuals from the ABCD Study with low-motion data at two consecutive time points. First, we calculated FC change scores between successive data collection time points (i.e., time 1 (*T*_*1*_) to time 2 (*T*_*2*_)), per participant, between 12 functional networks, using six different change score metrics, including simple change (ΔFC), number-line change (ΔFC+1), annualized percent change (APΔ, |APΔ|), and a reliable change index (RCT, |RCT|). Then, to understand the implications of each FC change score metric, we compared the results of the six metrics to each other using descriptive statistics. Due to evidence that connectomic measures comprising multiple connections are more reliable than individual connections (Yoo et al., 2019) and the prevalence of literature interpreting connectivity change in topological terms, we then calculated a number of graph theoretic metrics (i.e., clustering coefficient, betweenness centrality, network strength). To understand how each FC change score metric captures these ongoing topological changes, we calculated the dependence between each of the six FC change score metrics and changes in network-wise graph theoretic measures. Altogether, this work aims to inform future analytic choices for researchers hoping to study individual-centric FC changes, to further the study of individual differences in brain function by describing the impact of change score metrics on the interpretation of FC changes.

## Methods

### Participants

The longitudinal data used in this paper were collected as a part of the ongoing Adolescent Brain and Cognitive Development (ABCD) Study, and includes the annual 4.0 data release (http://dx.doi.org/10.15154/1523041). 11,880 children, ages 9 to 10 years (mean age = 9.49; 48% female), were enrolled in ABCD, a 10-year longitudinal study. Additional details on the ABCD Study sample and data collection are published elsewhere (Garavan et al., 2018; Volkow et al., 2018). Briefly, the ABCD Study’s sampling approach aimed to capture the nationwide sociodemographic diversity by largely recruiting participants at 21 study sites across the United States from elementary schools (private, public, and charter schools) (Garavan et al., 2018). The institutional review board and human research protections programs at the University of California San Diego approved all experimental and consent procedures, while each study site also obtained approval from their local institutional review board as well. Written assent was received from both the participants, as well as consent from their legal guardians, to participate in the study. ABCD Study exclusion criteria included age at enrollment (only youth ages 9.0 to 10.99 years included); no English fluency; MRI contraindications; history of traumatic brain injury or major neurological disorder; presence of any non-correctable sensory and/or motor impairments that would preclude the youth’s participation in study procedures; current intoxication at appointment; diagnosis of any DSM-I psychotic, autism spectrum, or substance use disorder; an intellectual disability reported by their caregiver; premature birth, very low birth weight, or perinatal complications; and caregiver knowledge at baseline of an impending move to an area beyond reasonable traveling distance to an ABCD Study site. Here, we use a subset of data from the ABCD Study, focused on resting-state functional magnetic resonance imaging (rsfMRI). Data used are from two time points which include assessments from baseline enrollment, at ages 9-11 years (i.e., *T*_1_), and year 2 follow-up, at ages 11-13 years (i.e., *T*_2_). Basic participant demographics and participants’ sex were all assessed at *T*_1_. For all following analyses, we limited our sample to youth with sufficient, high-quality rs-fMRI data at *T*_1_ and *T*_2_. Thus, we further excluded participants who: had incidental neurological findings evident in their scans, failed the ABCD imaging or FreeSurfer quality control procedures (Hagler et al., 2019), had 2-year follow-up imaging visits (i.e., *T*_*2*_) that took place after March 2020, exhibited greater than 0.5mm average head motion (i.e., framewise displacement) across rs-fMRI scans, and had fewer than 10 minutes of low-motion data (i.e., framewise displacement < 0.3 mm) at either time point (i.e., *T*_1_, *T*_2_). Of the 11,876 participants enrolled in the ABCD Study, only 3952 had high-quality, low-motion rs-fMRI data collected at both *T*_1_ and *T*_2_. Of those, only This resulted in an analytic sample of *N* = 2719 individuals. **Table 1** describes the final sample characteristics for the current study.

**Table 1.**
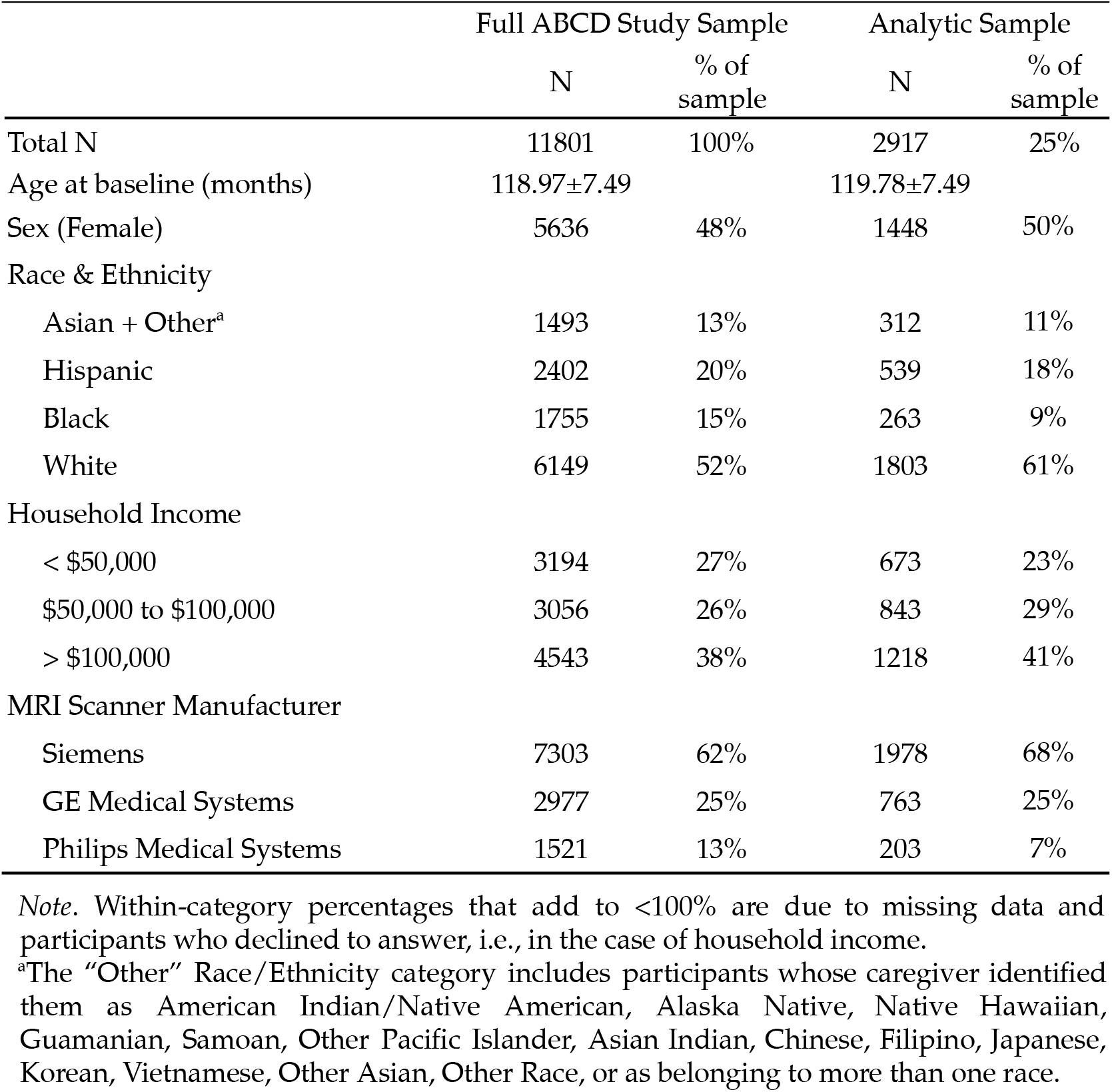
Sample Demographics.

### Neuroimaging Data

#### Structural MRI: Acquisition, Processing, and Quality Control

A harmonized data protocol was implemented across sites with either a Siemens, Phillips, or GE 3T MRI scanner. To reduce motion distortion, motion compliance training, as well as real-time, prospective motion correction was used. For more information on the scanning protocol, please see Casey et al. (2018). T1w images were acquired using a magnetization-prepared rapid acquisition gradient echo (MPRAGE) sequence and T2w images were obtained with fast spin echo sequence with variable flip angle (Casey et al., 2018). Both consist of 176 slices with 1 mm 3 isotropic resolution. Subjects were instructed to keep their eyes open and focused on a crosshair while twenty cumulative minutes of resting-state data was collected across two sets of two five-minute acquisition periods. These protocols increase the probability of collecting sufficient data with low motion per ABCD’s standards (>12.5 minutes of data with framewise displacement (FD) < 0.2 mm) (Power et al. 2014). Resting state scans were taken using an echo-planar imaging sequence in the axial plane. The following parameters were enforced for resting state scans: TR = 800 ms, TE = 30 ms, flip angle = 90°, voxel size = 2.4 mm^3^, 60 slices. Image processing steps have been previously described by Hagler and colleagues (2019).

Using the Gordon cortical network parcellation, functional connectivity (FC) between large-scale brain networks was previously estimated by calculating pairwise product-moment correlations of blood-oxygen-level-dependent (BOLD) signals between brain regions. These correlations were Fisher-transformed to z-scores and then averaged within each network defined by the Gordon parcellation (Hagler et al., 2019).

### Analyses

Prior to data analysis, a pre-registered analysis plan was registered with the <u>Open Science Framework (OSF)</u>. Outlier participants at each time point were identified using random forests as implemented in scikit-learn’s IsolationForest. Briefly, this approach isolates outlier participants by recursively dividing the sample based on values of a randomly selected variable (i.e., functional connection). Participants with outlier values on multiple FC variables end up isolated from the group relatively early in this recursive “tree”, thus identifying them as multivariate outliers. These outliers were removed from the sample prior to data analysis.

#### Functional connectivity (FC) Change Scores

To mitigate the impact of hardware- and motion-induced noise, MRI scanner serial number and head motion (i.e., average framewise displacement, FD) were regressed out from point estimates (i.e., FC estimates at T_1_, T_2_) prior to calculating FC change scores, using linear regression in pingouin (v0.5.1, pingouin-stats.org; Vallat, 2018). Then, four change scores metrics were calculated from the residuals of these models.

Sign changes for each functional connection were determined by assessing whether connectivity values were positive (i.e., FC > 0) or negative (i.e., FC < 0) at each *T*_1_ and *T*_2_. Sign changes indicate that the nature of the connectivity between networks (i.e., cooperating if FC > 0, opposing if FC < 0) changes between *T*_1_ and *T*_2_. Together with information regarding changes in the magnitude of connectivity, the sign changes help provide a more complete picture of information sharing (i.e., functional connectivity) between brain networks.

#### Simple change score, ΔFC

To illustrate its drawbacks, we include a simple measure of change, defined as the difference between residualized FC estimates at each data collection time point, scaled by the elapsed time between *T*_*1*_ and *T*_*2*_

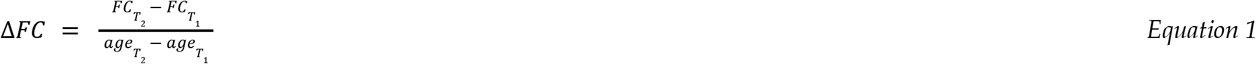

#### Number-line change score, ΔFC+1

For another simple estimate, we estimated the number-line change in FC, or the mathematical distance from FC at *T*_1_ and at *T*_2_ “along the number line” (i.e., disregarding zero and conflating the magnitude and nature of FC). To do so, we added 1 to FC estimates at each data collection time point to maintain the mathematical distance between them, then calculated the difference and scaled by the elapsed time between *T*_*1*_ and *T*_*2*_ (**ΔFC+1**; Equation 2).

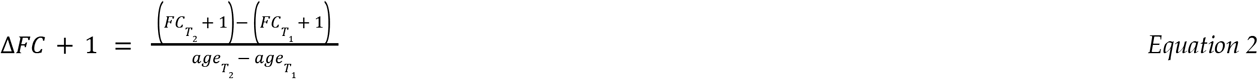

#### Annualized percent change score, APΔ

Annualized percent change is calculated as the difference between FC estimates at each data collection time point, scaled by the average of the two, over the elapsed time between *T*_*1*_ and *T*_*2*_. This method has been previously used in the developmental neuroimaging literature to estimate changes in measures of brain morphology, microarchitecture, and function (Mills et al., 2021; Bottenhorn et al., 2023). Here, we calculated the annualized percent change in *z*-scored FC estimates (**APΔ**; Equation 3) and in the absolute values thereof (|**APΔ**|).

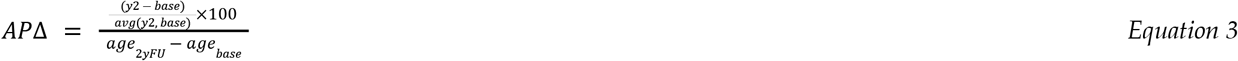

#### Reliable change over time score, RCT

Reliable change over time is an adaptation of the reliable change index (RCI) that incorporates the elapsed time between *T*_*1*_ and *T*_*2*_. While RCI is more commonly used to estimate clinical effects over a treatment period, it presents a method by which to incorporate different magnitudes of variance at each time point (Maassen, 2004) (Maassen, 2004). Here, we calculated the reliable change over time in *z*-scored FC estimates (**RCT**; Equation 4) and in the absolute values thereof (|**RCT**|).

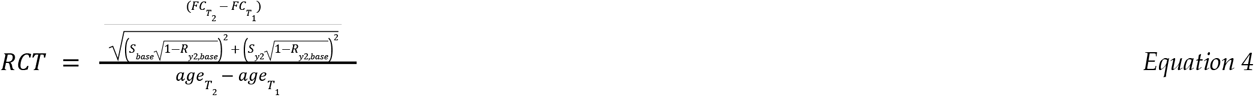

For each change score metric described above, we computed descriptive statistics (i.e., mean, standard deviation, range) for all connections of each network, for each network’s within-network connectivity, and for each network’s between-network connectivity.

#### Topology Changes

Graph theory metrics were calculated from residualized point FC estimates (i.e., *z*-scored FC at T_1_ and *T*_*2*_) for multivariate assessments of topology change for each network. These metrics included network strength, *k*_*i*_ (Equation 5), clustering coefficient, *C*_*i*_ (Equation 6), and betweenness centrality, *b*_*i*_ (Equation 7). Before calculating graph theory metrics, FC matrices were sparsified by calculating the Triangulated Maximally Filtered Graph. This approach is a topology-based alternative to thresholding, which has been shown to better identify reliable individual differences in functional brain networks (Jiang et al., 2023).

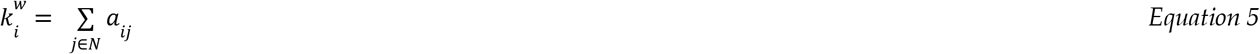

In calculating network strength, *k*_*i*_, using Equation 5, *a*_*ij*_ indicates connectivity weight (i.e., product-moment correlation) between nodes (i.e., networks) *i* and *j, N* is the set of all networks (Rubinov & Sporns, 2010). Here, network strength estimates the total connectedness of each network with the rest of the brain.

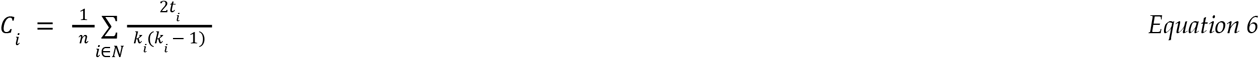

In calculating clustering coefficient, *C*_*i*_, using Equation 6, *n* indicates the number of networks, *t*_*i*_ is the number of triangles around network *i* (Rubinov & Sporns, 2010). Clustering coefficient is a measure of segregation, estimating how many of a network’s neighbors (i.e., directly connected networks) are connected to each other and, thus, the *clustering* around each network.

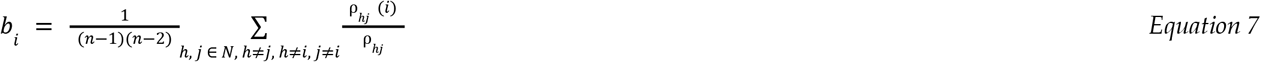

In calculating betweenness centrality, *b*_*i*_, using Equation 7, *ρ*_*hj*_ is the number of shortest paths between networks *h* and *j, ρ*_*hj*_*(i)* is the number of shortest paths between networks *h* and *j* that pass through network *i* (Rubinov & Sporns, 2010). Betweenness centrality is a measure of how often a network appears on the shortest path between other networks, indicating how integral a network is to the flow of information across the brain.

Changes in graph theory metrics were assessed, on an individual level, as a change over time (i.e., of the general form shown in Equation 8).

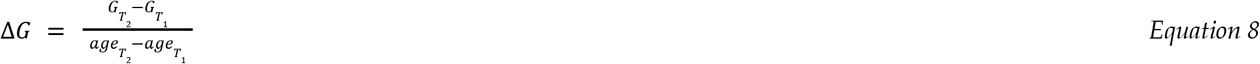

In Equation 5, *ΔG* represents the difference in each topological measure calculated as a difference in *G*_*Tx*_, the topological measure, at *T*_*1*_ and *T*_*2*_, scaled by the elapsed time between *T*_*1*_ and *T*_*2*_.

#### Correspondence between functional connectivity and topology changes

Then, we assessed correspondence between the changes in each topological measure, per network, and each FC change metric for that network’s connections using Spearman rank correlation as implemented in scipy (v1.10.1; (Abraham et al., 2014; Pedregosa et al., 2011)). Briefly, rank correlation estimates were chosen because of their intuitive interpretation: greater FC changes corresponding with greater topology changes yields a higher, positive rank correlation. Likewise, researchers are more likely to interpret greater increases in FC as “increased integration”; greater decreases, as “increased segregation”. Statistical significance each Spearman correlation estimate was determined via comparison with a null distribution simulated from randomly shu?ing the change metrics over 1000 iterations.

#### Correction for multiple comparisons

Significance was corrected for the number of effective comparisons across functional connections, assessed at *α*_*adjusted*_ < 0.05 (Li & Ji, 2005; Šidák, 1967). All 78 unique FC values (i.e., lower triangle of symmetric FC matrix) and the eigenvalues of a matrix of pairwise product-moment correlations between FC measures are used with this approach to determine the number of effective comparisons to correct for, accounting for dependence between measures. This approach revealed 43.76 “effective” comparisons across all 78 functional connections, adjusting *α* < to *α*_*adjusted*_ < 0.00117. This corrected threshold was used to determine significance for all reported *p*-values.

## Results

### FC change scores

Distributions of change estimates by each FC change score metric varied considerably (Figure 1), as did their average values for each connection (Figure 2). For some networks, change score distributions varied with respect to sign changes between time points, reflecting changes both in *magnitude* and *nature* of connectivity (Figure 1A).

**Figure 1.**
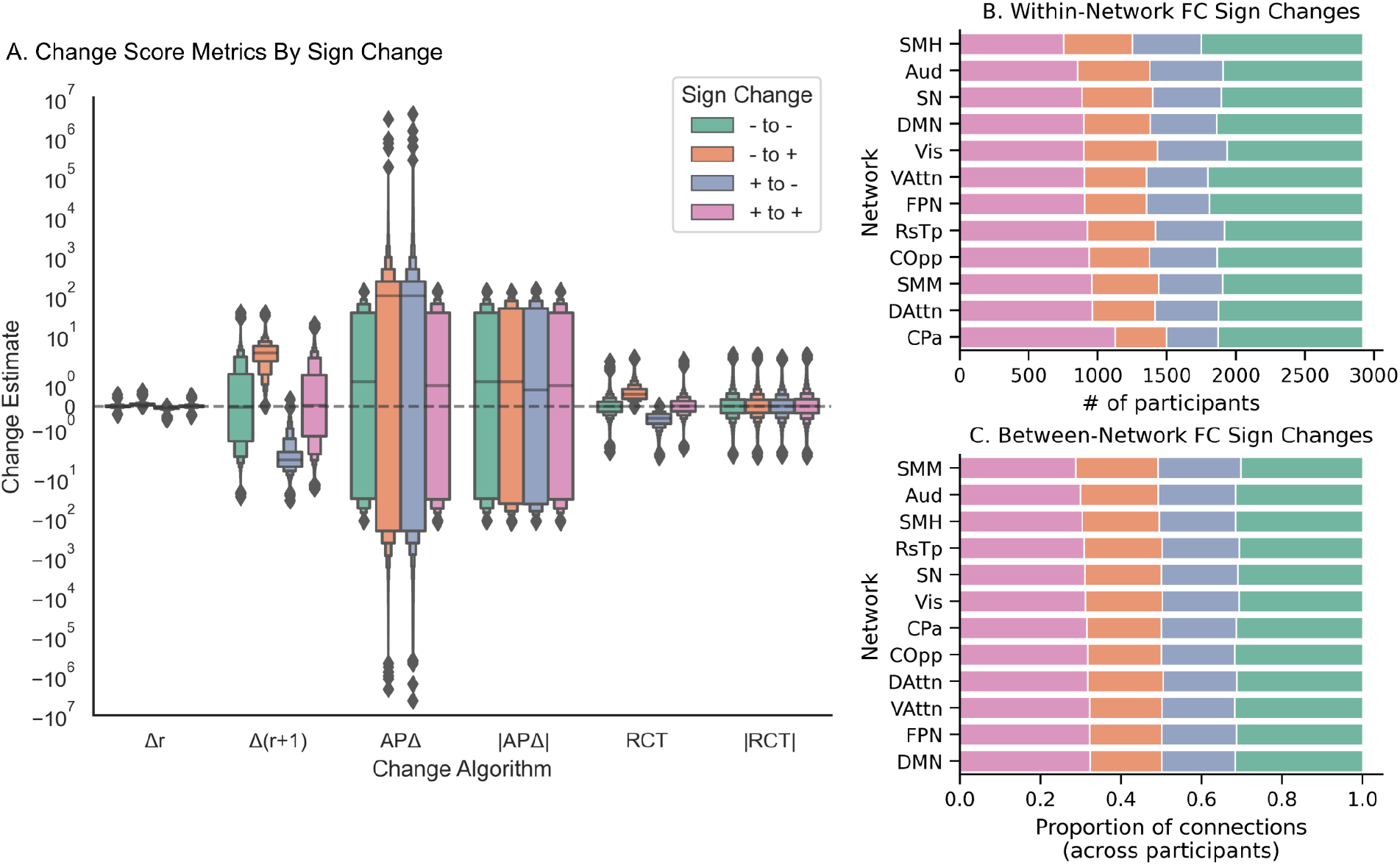
Estimates of change magnitudes across FC change scores, separated by the sign of the FC (i.e., negative, positive, and/or change in direction). (A) Different connectivity change metrics are presented on the *x*-axis. Distributions of individual change estimates across all networks are represented on the *y*-axis (log-scaled), separated and color-coded based on the FC sign (i.e., negative, positive, or change in direction from time 1 (*T*_*1*_) to time 2 (*T*_*2*_). (B) Distribution of within-network FC signs between *T*_*1*_ and *T*_*2*_. (C) Distribution of between-network FC signs between *T*_*1*_ and *T*_*2*_ per network.

**Figure 2.**
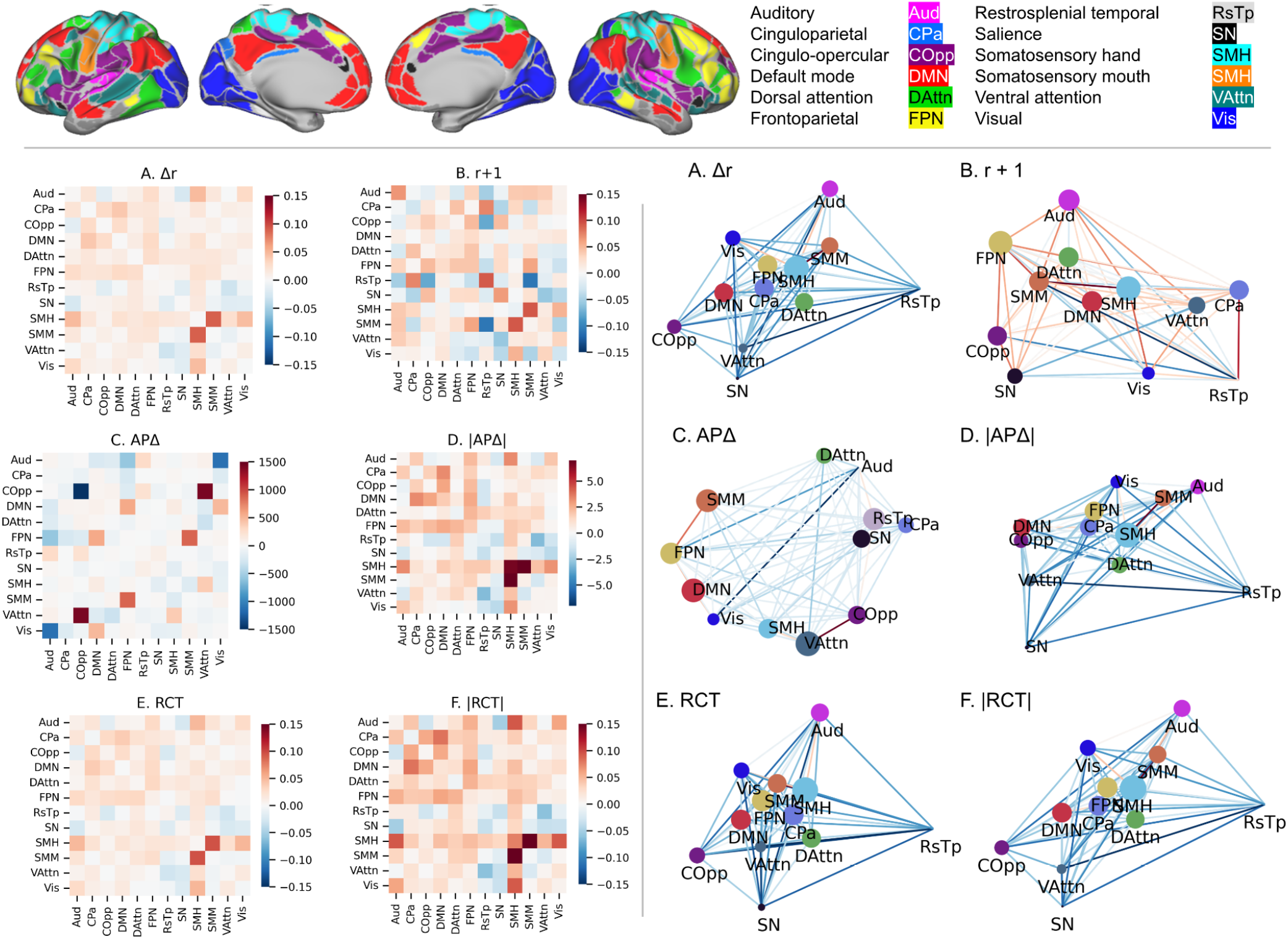
Average change scores across participants for each metric visualized by heatmaps and spring-embedded graphs. (Top): Functional brain networks analyzed here, as defined by Gordon et al. (2017) and their abbreviations (color-coded to match brain visualization and spring-embedded graph nodes). Matrices of average FC change score per metric (left) and spring-embedded graphs with edges weighted by their average value across participants (bottom): (A) simple change, (B) number-line change, (C) annualized percent change, (D) annualized change in absolute values, (E) reliable change over time, (F) reliable change in absolute values over time. Note that C and D are on different scales than the other plots, as annualized percent (absolute) change score means were orders of magnitude larger than those of the other change scores. Abbreviations: auditory network, Aud; cingulo-parietal network, CPa; cingulo-opercular network, COpp; default mode network, DMN; dorsal attention network, DAttn; fronto-parietal network, FPN; retrosplenial-temporal network, RsTp; salience network, SN; sensorimotor hand network, SMH; sensorimotor mouth network, SMM; ventral attention network, VAttn; visual network, Vis.

However, the majority of connections (both within- and between-networks) did not change sign between time points in most participants, i.e., cooperating connections remained cooperating (*r* > 0) and opposing connections remained opposing (*r* < 0) (Figure 1B). On average, across participants, estimates of simple change (Δr) were very near zero across connections, regardless of and did not differ between sign changes (Figures 1A, 2A top), while estimates of number line change (Δr + 1) were an order of magnitude larger (i.e., 10^0^ vs. 10^−2^) and reflected negative-to-positive and positive-to-negative sign changes as positive and negative values, respectively (Figures 1A, 2B top). On the other hand, estimates of annualized percent change (APΔ) were several orders of magnitude greater than both Δr and Δr + 1 (Figures 1A, 2C top), as were annualized percent absolute change (|APΔ|), though to a lesser extent (Figures 1A, 2D top), and neither distinguished between sign changes. Moreover, |APΔ| were more positive, on average, than Δr, Δr + 1, or APΔ. Finally, estimates of reliable change over time (RCT) and reliable absolute change over time (|RCT|) were smaller than Δr + 1, APΔ, and |APΔ|, on average (Figure 1A; Figure 2E & F, top), and while RCT reflected each negative-to-positive and positive-to-negative sign changes in the sign of its estimates, |RCT| did not (Figure 1A), for obvious, mathematical reasons.

#### Simple Change

Simple change (*Δr*) had the lowest and most consistent means and standard deviations across all change metrics, very near zero (Supplemental Table 1; Figure 2A, top). The greatest mean increases were within the auditory and retrosplenial-temporal networks (both *Δr*_*mean*_ = 0.0013), which can be interpreted as an average increase in correlation (i.e., connectivity) of 0.0013, while the greatest decreases were within the cinguloparietal network (*Δr*_*mean*_ = −0.0009). Within-network connectivity of the cinguloparietal network also demonstrated the greatest variability in simple change (*Δr*_*SD*_ = 0.148), while within-network connectivity of the frontoparietal network demonstrated the least variability in simple change (*Δr*_*SD*_ = 0.051).

#### Number Line Change

Overall, *Δr*+1 showed much greater variability than *Δr* (Supplemental Table 1; Figure 1A). Again, connectivity within retrosplenial temporal and auditory networks increased the most, per number-line change (*Δr+1*_*mean*_ = 0.0925; *Δr+1*_*mean*_ = 0.0667; Figure 2B), which can be interpreted as an average increase in *cooperativeness* of connectivity of 0.0925. Within-visual network connectivity had the greatest decreases (*Δr+1*_*mean*_ = −0.0302), which can be interpreted as an average increase in *oppositional* connectivity of 0.0302. Again, within-cinguloparietal network connectivity showed the greatest variability in number-line change (*Δr+1*_*SD*_ = 7.804; Supplemental Figure 1), while within-frontoparietal network connectivity showed the least variability (*Δr+1*_*SD*_ = 2.566).

#### Annualized Percent Change

Estimates of APΔ were the greatest and most variable, compared to other change score metrics, while |APΔ|were the second most variable (Supplemental Table 1; Figure 1A). Between-network connectivity of the cingulo-opercular network showed the greatest positive annual percent change (APΔ_*mean*_ = 112.83; Figure 2C), which can be interpreted as a ~113% mean increase in connectivity between cingulo-opercular and other networks. However, within-cingulo-opercular connectivity showed the greatest negative annual percentage change (APΔ_*mean*_ = −1478.00), which can be interpreted as a 1478% mean decrease in connectivity within cingulo-opercular. Within-cingulo-opercular also demonstrated the greatest variability in annual percent change (APΔ_*SD*_ = 80865; Supplemental Figure 1), while within-network connectivity of the salience network demonstrated the least variability (APΔ_*SD*_ = 1282.55).

Within-network connectivity of the somatosensory hand network demonstrated the greatest positive annual percent change in absolute values (|APΔ|_*mean*_ = 6.7208; Figure 2D), which can be interpreted as a ~6.7% mean increase in connectivity, regardless of the cooperative or opposing nature of the connections. Within-retrosplenial-temporal demonstrated the greatest negative annual percent change in absolute values (|APΔ|_*mean*_ = −0.6483), which can be interpreted as a ~0.65% mean decrease in connectivity, regardless of the cooperative or opposing nature of the connection. Between-network connectivity of the somatosensory mouth network demonstrated the greatest variability in annual percent change in absolute values (|APΔ|_*SD*_ = 52.076; Supplemental Figure 1); within-cinguloparietal connectivity, the least variability (|APΔ|_*SD*_ = 46.718).

#### Reliable Change Over Time

Second only to Δ*r*, reliable change over time (RCT) and reliable change in absolute values over time (|RCT|) demonstrated the lowest standard deviations across change score metrics (Supplemental Table 1; Figure 1A). Both change score metrics demonstrated mean values similar in magnitude to Δ*r* and Δ*r* + 1. Within-auditory network connectivity demonstrated the greatest increase in RCT (RCT_*mean*_ = 0.0091; Figure 2E), which can be interpreted as an increase in connectivity that is about 1% of the magnitude of the standard error in connectivity across the sample. Within-cinguloparietal network connectivity demonstrated the greatest decrease in RCT (RCT_*mean*_ = −0.0036). Due to the nature of the RCT calculation, variability was largely the same across connections (RCT_*SD*_ = 0.504-0.508; Supplemental Figure 1).

Within-visual network connectivity demonstrated the greatest increase in reliable change in absolute values over time (|RCT|_mean_ = 0.0506; Figure 2F), which can be interpreted as an increase in connectivity that is about 5% of the magnitude of the standard error in connectivity across the sample. Average auditory network connectivity demonstrated the greatest decrease in RCT (|RCT|_mean_ = −0.0118). As with RCT, variability was largely the same across connections (|RCT|_*SD*_ = 0.505-0.507; Supplemental Figure 1).

### Topology Changes

Even before comparisons with graph theoretic measures, there is some suggestion that ΔFC metrics imply differing changes in topology (Figure 2A-F, right). Spring embedded graphs of average change scores based on each metric are shown in the bottom of Figure 2. In these graphs, the distance between nodes (i.e., networks) represents the magnitude of change such that nodes becoming more connected are closer together, while nodes becoming less connected are further apart. Moreover, the size of the circle representing a node is proportional to its change in overall connectedness (i.e., strength) such that larger circles indicate increased average connectivity of that network, while edges are color-coded to match their values on the above matrices (Figure 2, left). Averaged across participants, most change scores imply that the somatosensory (hand) network is becoming *more* connected and *more* central (Figure 2, right) while RsTp is becoming *less* so, but APΔ shows the opposite trends (Figure 2C, right). Further, Δ*r*, |APΔ|, RCT, and |RCT| suggest that the frontoparietal network is becoming more connected, on the whole, while Δ*r* + 1 and APΔ suggest it is becoming less connected. However, all six change score metrics agree that Aud is becoming less connected over time.

When calculating network-level topological differences between time points (Equation 8), distinct trends emerged for clustering coefficient and betweenness centrality (Figure 3, Table 2). Half of the networks’ clustering coefficients decreased, while the other half increased between the two time points. In contrast, changes in betweenness centrality were much smaller: the majority of networks showed little to no change in betweenness centrality estimates between timepoints.

**Table 2.**
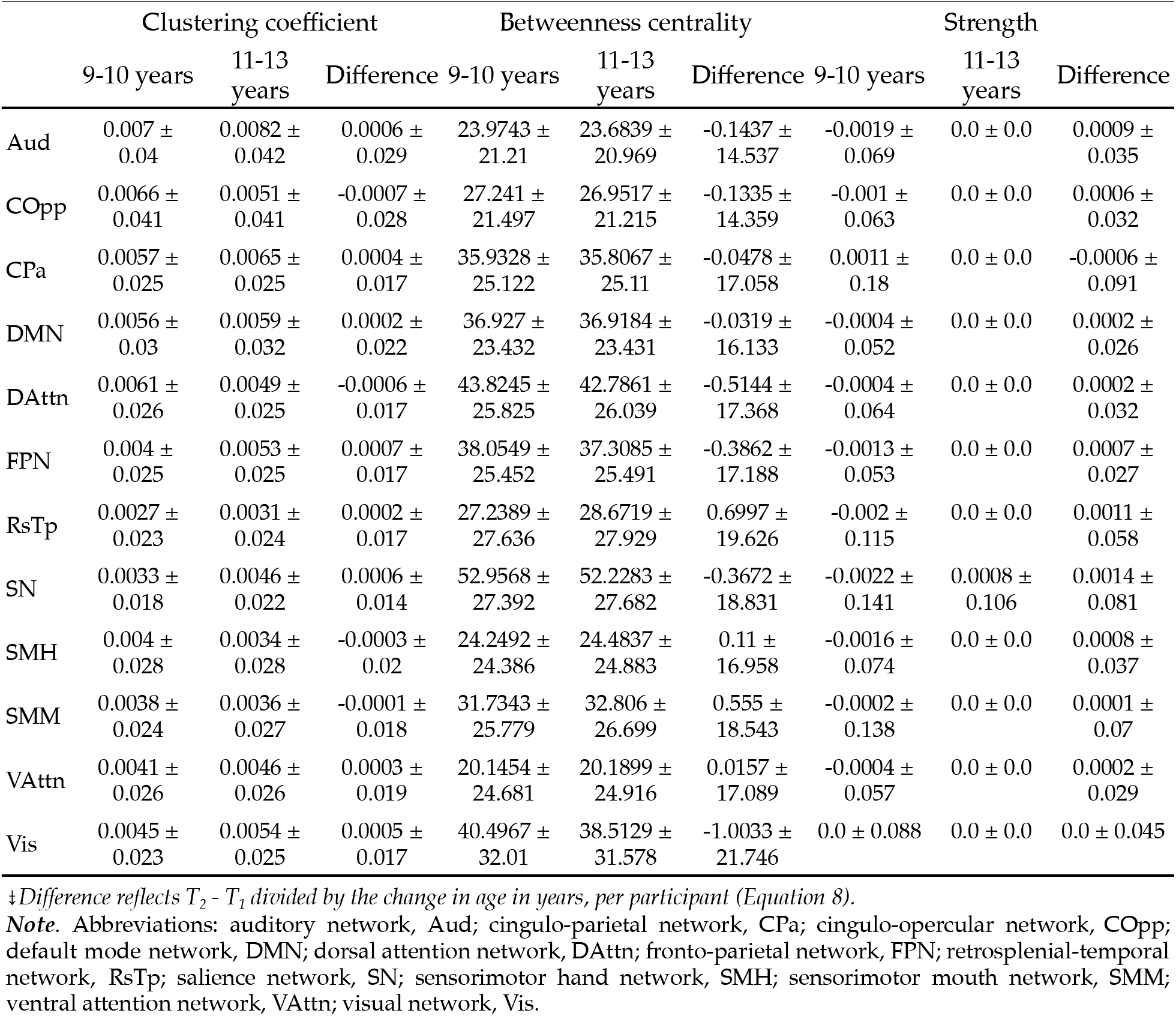
Mean and standard deviation of graph theoretic measures at each data collection time point, and the difference between T_1_ and T_2_, scaled by time elapsed.

**Figure 3.**
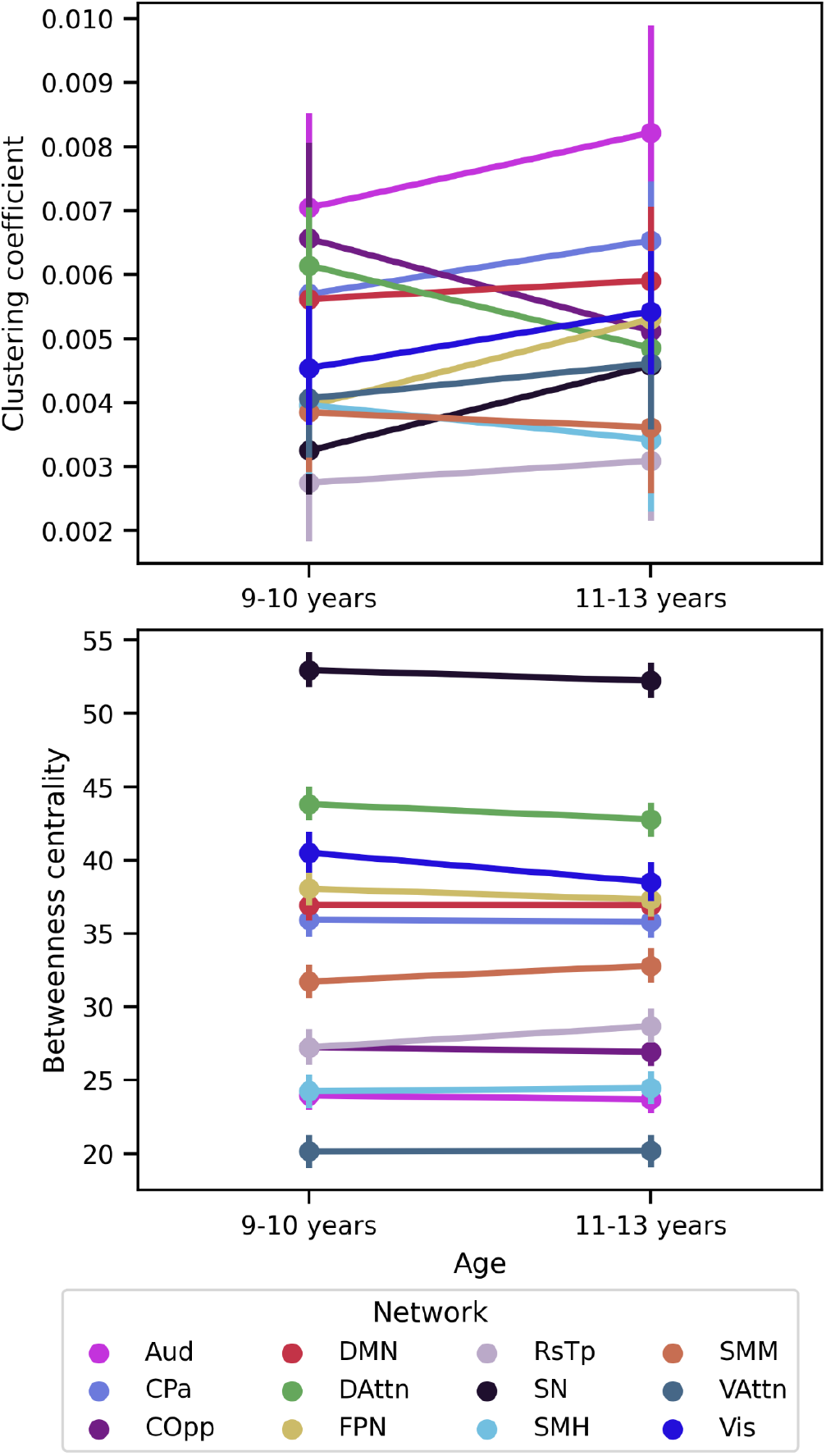
Local functional connectome properties at ages 9-10 and 11-13 years. Data collection time point is shown on the *x*-axes and graph theoretic values per network (across participants: mean as the point ± standard deviation as vertical lines) are indicated by the *y*-axes for clustering coefficient (*Equation 6*) on the top and betweenness centrality (*Equation 7*) on the bottom. Network strength is not included because the average strength (i.e., sum of connectivity) for each network at each time point is zero, due to confound regression. Abbreviations: auditory network, Aud; cingulo-parietal network, CPa; cingulo-opercular network, COpp; default mode network, DMN; dorsal attention network, DAttn; fronto-parietal network, FPN; retrosplenial-temporal network, RsTp; salience network, SN; sensorimotor hand network, SMH; sensorimotor mouth network, SMM; ventral attention network, VAttn; visual network, Vis. See Table 2 for mean and standard deviation of values at each time point and the difference between time points.

### Correspondence between functional connectivity and topology changes

Network-average dependence between FC change metrics and topological changes, estimated from Spearman rank correlations, were small (−0.075 < *r*_*S*_ < 0.15). However, there were different trends (i.e., relative, across networks) in correspondence between FC change metrics for each aspect of topological change (Figure 4; Supplementary Table 1). While RCT exhibited the most significant correlations with betweenness centrality changes (Figure 4A), across networks, RCT, Δr, and Δr+1 all demonstrated the most significant correlations with changes in clustering coefficient (Figure 4B) and network strength (Figure 4C).

**Figure 4.**
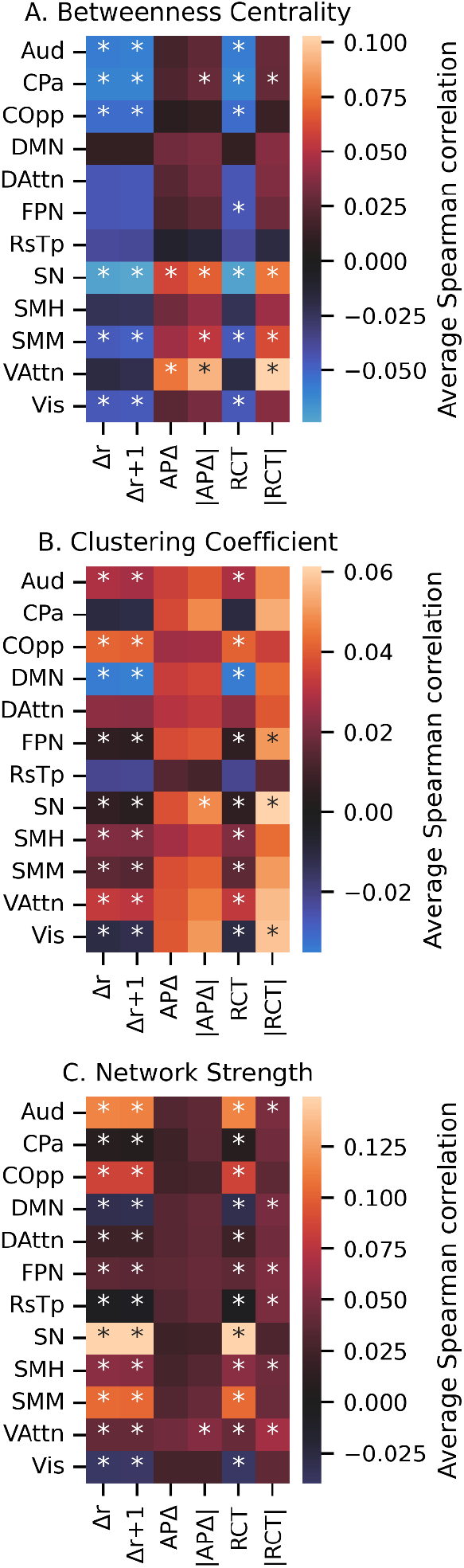
Average Spearman correlation between each FC change metric and differences in graph theory measures as calculated at each wave separately. Asterisks (*) denote networks for which the majority of connections exhibit significant rank correlation with the topological measure: how FC changes capture differences in (A) betweenness centrality, (B) clustering coefficient, and (C) network strength between waves of data collection. Abbreviations: auditory network, Aud; cingulo-parietal network, CPa; cingulo-opercular network, COpp; default mode network, DMN; dorsal attention network, DAttn; fronto-parietal network, FPN; retrosplenial-temporal network, RsTp; salience network, SN; sensorimotor hand network, SMH; sensorimotor mouth network, SMM; ventral attention network, VAttn; visual network, Vis.

## Discussion

The current study compares several different metrics for assessing changes in functional connectivity (FC) between two data collection time points, using data from over 2,700 youth at ages 9-10 years (i.e., *T*_*1*_) and again at 11-13 years (i.e., *T*_*2*_). In doing so, we aimed to reconcile that FC estimates, reflecting correlations of blood-oxygen-level-dependent time courses contain two unique pieces of information: the magnitude (i.e., shared information between regions) and the direction (i.e., sign reflecting cooperation (i.e., *r* > 0) or opposition (i.e., *r* < 0)) of the connectivity. Thus, studies that only have two repeated measures face a unique challenge: estimating FC changes, while also preserving both pieces of information. Our findings highlight how the choice of the resting state change metric can impact both the estimation of functional connectivity changes and their interpretation. Researchers should consider these mathematical and conceptual differences when deciding how to estimate FC changes, as they could lead to very different results and interpretations.

These results strongly suggest that FC change scores not only vary mathematically, but also in the aspects of FC that they capture. While APΔ has been previously used in the developmental neuroimaging literature to discuss early adolescent brain changes (Bottenhorn et al., 2023; Mills et al., 2021), our findings show it is more suited to describing structural changes than correlation-based FC, which carry inherent magnitude and direction information in a single score. Estimates of APΔ ranged were orders of magnitude larger than those of other change metrics, ranging from −10^5^ to +10^5^, which would boil down to more than a ten thousand percent change per year in the brain’s functional connectivity, which seems unlikely. When using the absolute values of annual percent change (i.e., |APΔ|), the range of ΔFC estimates was a little narrower, but still indicated nearly a 100% change per year in some networks. These large positive and negative APΔ estimates are driven by near-zero denominators caused by averaging positive and negative connectivity estimates of similar absolute values. Further, both APΔ and |APΔ|demonstrated on average more positive changes between networks, regardless of sign changes between time points. This indicates that, while they may capture changes in the amount of information shared between regions or networks (hereafter, “nodes”) they poorly differentiate the nature of connections (i.e., cooperative vs. opposing brain activation). On the other end of the spectrum, straight-forward simple change (Δr) estimates were near zero with little variation, but this metric did a better job preserving sign changes between the two time points, suggesting cooperative changes and opposing changes are largely captured. Number-line change (i.e., Δr+1), likewise, had a narrower range but on the order of ±10^1^, and similarly differentiated sign changes in FC. Finally, reliable changes over time (i.e., RCT) and reliable absolute changes over time (|RCT|) were both within ±10^1^ and, while RCT differentiated between FC sign changes, |RCT| did not. To summarize, APΔ and |APΔ| present potentially unrealistic estimates of ΔFC, while Δr, Δr+1, RCT, and |RCT| may more faithfully capture the magnitude of changes. Additionally, Δr, Δr+1, and RCT preserve information about the nature of connectivity changes, while APΔ, |APΔ|, and |RCT| did not. Thus, of all metrics examined, Δr, Δr+1, and RCT were deemed better in their ability to estimate both magnitude and sign of changes in resting state functional connectivity.

Many studies that assess changes in functional connectivity frame their findings in terms of “integration” and “segregation”, especially in the developmental neuroimaging literature (Cohen & D’Esposito, 2016; Fair et al., 2007; Grayson & Fair, 2017). These concepts provide intuitive descriptions of connectivity increasing and decreasing the amount of shared information and cooperation between nodes. In addition to these informal interpretations, network science and graph theory offer formalized mathematical definitions of “integration” and “segregation”, and other properties like “centrality”. In network science, segregation indicates functional separation between nodes (here, networks) for specialized processing, while integration refers to the combination of information from such nodes’ separated, specialized processing (Rubinov & Sporns, 2010). Centrality also suggests a role in functional integration by reflecting how important a node is to the rest of the network or, in this case, the rest of the brain (Rubinov & Sporns, 2010). By using graph theoretic metrics, we aimed to further assess how each of the functional connectivity change metrics can capture higher-order topological features. In general, greater ΔFC across metrics was associated with greater clustering coefficient and network strength. The few exceptions to this trend come from the default mode network, in which ΔFC change was inversely associated with changes in clustering coefficient and network strength, although weakly. We found that betweenness centrality was best captured by RCT, based on average Spearman correlation, suggesting that RCT can capture changes in a network’s centrality, importance, and ability to integrate information. On the other hand, |RCT|, APΔ, and |APΔ| all suggested opposite associations of betweenness centrality and ΔFC, compared to the other metrics: |RCT|, APΔ, and |APΔ| suggest that increased FC corresponds with greater betweenness centrality, while Δr, Δr+1, and RCT all suggest that increasing FC corresponds with diminished betweenness centrality. While clustering coefficient and network strength were best captured by the ΔFC metrics that were also found to more accurately describe magnitude and sign changes, including Δr, Δr+1, and RCT. This means that these three metrics can accurately reflect changes in segregation and in a network’s overall connectedness. Aside from a few contradictory betweenness centrality associations, neither APΔ, nor |APΔ| captured topological changes with any consistency. When assessing network-specific patterns, we found no ΔFC accurately captured betweenness centrality changes of the default mode, dorsal attention, retrosplenial temporal, or somatosensory (hand) networks, or clustering coefficient changes in cinguloparietal, dorsal attention, or retrosplenial temporal networks. In contrast, APΔ, |APΔ|, and |RCT| findings showed the somatosensory hand network becoming more connected while the ventral attention network became more disconnected. These additional topology analyses show how different ways of estimating individual differences in resting state functional connectivity can yield drastic differences in interpretations, in both globally and network-specific ways.

### Recommendations

Our work suggests that researchers should consider several of the mathematical and topological factors when choosing how to estimate individual-level functional connectivity changes between two time points.

1. Mathematically, APΔ and |APΔ| appear to vastly overestimate individual differences in functional connectivity changes per year (i.e., suggesting FC changes over 1000%), while Δr likely underestimates these findings (i.e., suggesting correlation changes around *r* = 0.01). Thus, APΔ should not be used to estimate ΔFC. Researchers should, instead, consider Δr+1, RCT, and |RCT|.
2. Conceptually, Δr, Δr+1, and RCT capture the change in the magnitude of shared information between nodes, and more accurately capture changes from an opposing (i.e., *r* < 0) to cooperative (i.e., *r* > 0) relationships as positive changes and changes from cooperative to opposing relationships as negative. In contrast, APΔ, |APΔ|, and |RCT| did not distinguish between the two. Researchers should consider the conceptual implications of sign changes in the context of their study when choosing a functional connectivity change score metric.
3. Topologically, the choice of functional connectivity change score metric can further complicate high-level interpretations. While centrality changes were not particularly well-represented by ΔFC metrics for many networks, changes in segregation and node connectedness (i.e., clustering coefficient and strength, respectively) were represented well by Δr, Δr+1, and RCT.

Further, the choice of functional connectivity metric should depend, too, on the sample and study design. For example, in studies of child development, and of some disease states, head motion may be a major source of measurement error that varies between time points (e.g., with age, disease severity) and can, thus, confound estimates of changes in functional connectivity. Even after censoring high-motion volumes and regressing out motion time series, the effects of head motion persist and can impart distance-dependent artifacts in functional connectivity estimates (Power et al., 2012). In such cases, RCT might especially be a good choice, as it scales change by the standard errors, allowing for unique variances at each time point. This study has focused entirely on change between two time points, due to the utility and preponderance of pre/post designs and the high costs of acquiring magnetic resonance imaging data that compound with additional data points. However, it is worth noting that such designs are unable to separate measurement error from true change and individual differences therein. However, if researchers have multiple resting-state fMRI runs at each time point, estimating change scores separately for each run, and then assembling a composite functional change score measure may help account for measurement noise that is otherwise difficult to address with only two point estimates. Further, there is a considerable body of literature on test-retest reliability in fMRI and functional connectivity (Noble et al., 2019, 2021), which may help contextualize observed changes and provide an additional means by which to estimate a reliability coefficient. Lastly, while assessing regression to the mean is beyond the scope of this work, researchers using two-time-point designs should remain aware and conscious of the way extreme values may introduce such effects (Barnett et al., 2005).

## Conclusion

While comparing functional connectivity between two time points (e.g., across development, pre-vs. post-intervention or -injury) can provide useful insights into brain changes, the interpretation of such changes can differ considerably between methods for estimating change. This work profiles such methods and compares them to topology changes, in a sample of over 2,700 youth with repeated resting-state fMRI measurements.

## Acknowledgments

A special thank you to all the children and families for their participation in their ABCD Study.

Research described in this article was supported by the National Institutes of Health (MMH: NIEHS R01ES032295, R01ES031074; KLB: K99MH135075).

Data used in the preparation of this article were obtained from the Adolescent Brain Cognitive Development^SM^ (ABCD) Study (https://abcdstudy.org), held in the NIMH Data Archive (NDA). This is a multisite, longitudinal study designed to recruit more than 10,000 children age 9-10 and follow them over 10 years into early adulthood. The ABCD Study® is supported by the National Institutes of Health and additional federal partners under award numbers U01DA041048, U01DA050989, U01DA051016, U01DA041022, U01DA051018, U01DA051037, U01DA050987, U01DA041174, U01DA041106, U01DA041117, U01DA041028, U01DA041134, U01DA050988, U01DA051039, U01DA041156, U01DA041025, U01DA041120, U01DA051038, U01DA041148, U01DA041093, U01DA041089, U24DA041123, U24DA041147. A full list of supporters is available at https://abcdstudy.org/federal-partners.html. A listing of participating sites and a complete listing of the study investigators can be found at https://abcdstudy.org/consortium_members/. ABCD consortium investigators designed and implemented the study and/or provided data but did not necessarily participate in the analysis or writing of this report. This manuscript reflects the views of the authors and may not reflect the opinions or views of the NIH or ABCD consortium investigators. The ABCD data repository grows and changes over time. The ABCD data used in this report came from http://dx.doi.org/10.15154/z563-zd24.

## Author Contributions

Conceptualization: KLB, MMH

Data curation: KLB

Formal Analysis: KLB, JDC

Funding acquisition: MMH, KLB

Methodology: KLB, MMH, HA

Project administration: KLB, MMH

Resources: MMH

Software: KLB Supervision: MMH

Visualization: KLB, JDC

Writing – original draft: KLB, JDC, MMH

Writing – review & editing: KLB, JDC, MMH, HA

## References

Abraham, A., Pedregosa, F., Eickenberg, M., Gervais, P., Mueller, A., Kossaifi, J., Gramfort, A., Thirion, B., & Varoquaux, G. (2014). Machine learning for neuroimaging with scikit-learn. Frontiers in Neuroinformatics, 8, 14. 10.3389/fninf.2014.00014

Bandettini, P. A., Gonzalez-Castillo, J., Handwerker, D., Taylor, P., Chen, G., & Thomas, A. (2022). The challenge of BWAs: Unknown unknowns in feature space and variance. Med, 3(8), 526–531. 10.1016/j.medj.2022.07.002

Barnett, A. G., van der Pols, J. C., & Dobson, A. J. (2005). Regression to the mean: What it is and how to deal with it. International Journal of Epidemiology, 34(1), 215–220. 10.1093/ije/dyh299

Betzel, R. F., Byrge, L., He, Y., Goñi, J., Zuo, X.-N., & Sporns, O. (2014). Changes in structural and functional connectivity among resting-state networks across the human lifespan. NeuroImage, 102, 345–357. 10.1016/j.neuroimage.2014.07.067

Boly, M., Phillips, C., Tshibanda, L., Vanhaudenhuyse, A., Schabus, M., Dang-Vu, T. t., Moonen, G., Hustinx, R., Maquet, P., & Laureys, S. (2008). Intrinsic Brain Activity in Altered States of Consciousness. Annals of the New York Academy of Sciences, 1129(1), 119–129. 10.1196/annals.1417.015

Bottenhorn, K. L., Cardenas-Iniguez, C., Mills, K. L., Laird, A. R., & Herting, M. M. (2023). Profiling intra- and inter-individual differences in brain development across early adolescence. NeuroImage, 279, 120287. 10.1016/j.neuroimage.2023.120287

Bottenhorn, K. L., Salo, T., Jacobs, E. G., Pritschet, L., Taylor, C., Herting, M. M., & Laird, A. R. (2024). Idiosyncrasy and generalizability of contraceptive- and hormone-related functional connectomes across the menstrual cycle (2024.10.24.620112; p. 2024.10.24.620112). bioRxiv. 10.1101/2024.10.24.620112

Button, K. S., Ioannidis, J. P. A., Mokrysz, C., Nosek, B. A., Flint, J., Robinson, E. S. J., & Munafò, M. R. (2013). Power failure: Why small sample size undermines the reliability of neuroscience. Nature Reviews Neuroscience, 14(5), 365–376. 10.1038/nrn3475

Casey, B. J., Cannonier, T., Conley, M. I., Cohen, A. O., Barch, D. M., Heitzeg, M. M., Soules, M. E., Teslovich, T., Dellarco, D., Garavan, H., Orr, C. A., Wager, T. D., Banich, M. T., Speer, N. K., Sutherland, M. T., Riedel, M. C., Dick, A. S., Bjork, J. M., Thomas, K. M., … Dale, A. M. (2018). The Adolescent Brain Cognitive Development (ABCD) study: Imaging acquisition across 21 sites. Developmental Cognitive Neuroscience, 32, 43–54. 10.1016/J.DCN.2018.03.001

Chai, X. J., Castañón, A. N., Öngür, D., & Whitfield-Gabrieli, S. (2012). Anticorrelations in resting state networks without global signal regression. NeuroImage, 59(2), 1420–1428. 10.1016/j.neuroimage.2011.08.048

Chan, M. Y., Park, D. C., Savalia, N. K., Petersen, S. E., & Wig, G. S. (2014). Decreased segregation of brain systems across the healthy adult lifespan. Proceedings of the National Academy of Sciences, 111(46), E4997–E5006. 10.1073/pnas.1415122111

Cohen, J. R., & D’Esposito, M. (2016). The Segregation and Integration of Distinct Brain Networks and Their Relationship to Cognition. The Journal of Neuroscience, 36(48), 12083. 10.1523/JNEUROSCI.2965-15.2016

Fair, D. A., Dosenbach, N. U. F., Church, J. A., Cohen, A. L., Brahmbhatt, S., Miezin, F. M., Barch, D. M., Raichle, M. E., Petersen, S. E., & Schlaggar, B. L. (2007). Development of distinct control networks through segregation and integration. Proceedings of the National Academy of Sciences, 104(33), 13507–13512. 10.1073/pnas.0705843104

Garavan, H., Bartsch, H., Conway, K., Decastro, A., Goldstein, R. Z., Heeringa, S., Jernigan, T., Potter, A., Thompson, W., & Zahs, D. (2018). Recruiting the ABCD sample: Design considerations and procedures. Developmental Cognitive Neuroscience, 32, 16–22. 10.1016/j.dcn.2018.04.004

Gordon, E. M., Laumann, T. O., Gilmore, A. W., Newbold, D. J., Greene, D. J., Berg, J. J., Ortega, M., Hoyt-Drazen, C., Gratton, C., Sun, H., Hampton, J. M., Coalson, R. S., Nguyen, A. L., McDermott, K. B., Shimony, J. S., Snyder, A. Z., Schlaggar, B. L., Petersen, S. E., Nelson, S. M., & Dosenbach, N. U. F. (2017). Precision Functional Mapping of Individual Human Brains. Neuron, 95(4), 791-807.e7. 10.1016/j.neuron.2017.07.011

Gracia-Tabuenca, Z., Díaz-Patiño, J. C., Arelio-Ríos, I., Moreno-García, M. B., Barrios, F. A., & Alcauter, S. (2023). Development of the Functional Connectome Topology in Adolescence: Evidence from Topological Data Analysis. eNeuro, 10(2), ENEURO.0296-21.2022. 10.1523/ENEURO.0296-21.2022

Gratton, C., Laumann, T. O., Nielsen, A. N., Greene, D. J., Gordon, E. M., Gilmore, A. W., Nelson, S. M., Coalson, R. S., Snyder, A. Z., Schlaggar, B. L., Dosenbach, N. U. F., & Petersen, S. E. (2018). Functional Brain Networks Are Dominated by Stable Group and Individual Factors, Not Cognitive or Daily Variation. Neuron, 98(2), 439-452.e5. 10.1016/j.neuron.2018.03.035

Grayson, D. S., & Fair, D. A. (2017). Development of large-scale functional networks from birth to adulthood: A guide to the neuroimaging literature. NeuroImage, 160, 15–31. 10.1016/j.neuroimage.2017.01.079

Greicius, M. D., Krasnow, B., Reiss, A. L., & Menon, V. (2003). Functional connectivity in the resting brain: A network analysis of the default mode hypothesis. Proceedings of the National Academy of Sciences of the United States of America, 100(1), 253–258. 10.1073/pnas.0135058100

Hagler, D. J., Hatton, S., Cornejo, M. D., Makowski, C., Fair, D. A., Dick, A. S., Sutherland, M. T., Casey, B. J., Barch, D. M., Harms, M. P., Watts, R., Bjork, J. M., Garavan, H. P., Hilmer, L., Pung, C. J., Sicat, C. S., Kuperman, J., Bartsch, H., Xue, F., … Dale, A. M. (2019). Image processing and analysis methods for the Adolescent Brain Cognitive Development Study. NeuroImage, 202, 116091. 10.1016/j.neuroimage.2019.116091

Hlinkaa, J., Paluša, M., Vejmelkaa, M., Mantini, D., & Corbetta, M. (2011). Functional connectivity in resting-state fMRI: Is linear correlation sufficient? NeuroImage, 54(3), 2218–2225. 10.1016/j.neuroimage.2010.08.042

Jiang, C., He, Y., Betzel, R. F., Wang, Y.-S., Xing, X.-X., & Zuo, X.-N. (2023). Optimizing network neuroscience computation of individual differences in human spontaneous brain activity for test-retest reliability. Network Neuroscience, 7(3), 1080–1108. 10.1162/netn_a_00315

Laird, A. R. (2021). Large, open datasets for human connectomics research: Considerations for reproducible and responsible data use. NeuroImage, 244, 118579.

Li, J., & Ji, L. (2005). Adjusting multiple testing in multilocus analyses using the eigenvalues of a correlation matrix. Heredity, 95(3), 221–227. 10.1038/sj.hdy.6800717

Maassen, G. H. (2004). The standard error in the Jacobson and Truax Reliable Change Index: The classical approach to the assessment of reliable change. Journal of the International Neuropsychological Society, 10(6), 888–893. 10.1017/S1355617704106097

Marek, S., Tervo-Clemmens, B., Calabro, F. J., Montez, D. F., Kay, B. P., Hatoum, A. S., Donohue, M. R., Foran, W., Miller, R. L., Hendrickson, T. J., Malone, S. M., Kandala, S., Feczko, E., Miranda-Dominguez, O., Graham, A. M., Earl, E. A., Perrone, A. J., Cordova, M., Doyle, O., … Dosenbach, N. U. F. (2022). Reproducible brain-wide association studies require thousands of individuals. Nature, 603(7902), 654–660. 10.1038/s41586-022-04492-9

Mills, K. L., Siegmund, K. D., Tamnes, C. K., Ferschmann, L., Wierenga, L. M., Bos, M. G. N., Luna, B., Li, C., & Herting, M. M. (2021). Inter-individual variability in structural brain development from late childhood to young adulthood. NeuroImage, 242, 118450. 10.1016/j.neuroimage.2021.118450

Noble, S., Scheinost, D., & Constable, R. T. (2019). A decade of test-retest reliability of functional connectivity: A systematic review and meta-analysis. NeuroImage, 203, 116157. 10.1016/j.neuroimage.2019.116157

Noble, S., Scheinost, D., & Constable, R. T. (2021). A guide to the measurement and interpretation of fMRI test-retest reliability. Current Opinion in Behavioral Sciences, 40, 27–32. 10.1016/j.cobeha.2020.12.012

Pedregosa, F., Varoquaux, G., Gramfort, A., Michel, V., Thirion, B., Grisel, O., Blondel, M., Prettenhofer, P., Weiss, R., Dubourg, V., Vanderplas, J., Passos, A., Cournapeau, D., Brucher, M., Perrot, M., & Duchesnay, É. (2011). Scikit-learn: Machine Learning in Python. Journal of Machine Learning Research, 12(Oct), 2825–2830.

Poldrack, R. A., Laumann, T. O., Koyejo, O., Gregory, B., Hover, A., Chen, M.-Y., Gorgolewski, K. J., Luci, J., Joo, S. J., Boyd, R. L., Hunicke-Smith, S., Simpson, Z. B., Caven, T., Sochat, V., Shine, J. M., Gordon, E., Snyder, A. Z., Adeyemo, B., Petersen, S. E., … Mumford, J. A. (2015). Long-term neural and physiological phenotyping of a single human. Nature Communications, 6, 8885.

Power, J. D., Barnes, K. A., Snyder, A. Z., Schlaggar, B. L., & Petersen, S. E. (2012). Spurious but systematic correlations in functional connectivity MRI networks arise from subject motion. Neuroimage, 59(3), 2142–2154.

Pritschet, L., Santander, T., Taylor, C. M., Layher, E., Yu, S., Miller, M. B., Grafton, S. T., & Jacobs, E. G. (2020). Functional reorganization of brain networks across the human menstrual cycle. NeuroImage, 220, 117091.10.1016/j.neuroimage.2020.117091

Rubinov, M., & Sporns, O. (2010). Complex network measures of brain connectivity: Uses and interpretations. NeuroImage, 52(3), 1059–1069. 10.1016/j.neuroimage.2009.10.003

Saha, R., Saha, D. K., Rahaman, M. A., Fu, Z., Liu, J., & Calhoun, V. D. (2024). A Method to Estimate Longitudinal Change Patterns in Functional Network Connectivity of the Developing Brain Relevant to Psychiatric Problems, Cognition, and Age. Brain Connectivity, 14(2), 130–140. 10.1089/brain.2023.0040

Šidák, Z. (1967). Rectangular Confidence Regions for the Means of Multivariate Normal Distributions. Journal of the American Statistical Association, 62(318), 626–633. 10.1080/01621459.1967.10482935

Smith, S. M., Fox, P. T., Miller, K. L., Glahn, D. C., Fox, P. M., Mackay, C. E., Filippini, N., Watkins, K. E., Toro, R., Laird, A. R., & Beckmann, C. F. (2009). Correspondence of the brain’s functional architecture during activation and rest. Proceedings of the National Academy of Sciences of the United States of America, 106(31), 13040–13045. 10.1073/pnas.0905267106

Vij, S. G., Nomi, J. S., Dajani, D. R., & Uddin, L. Q. (2018). Evolution of spatial and temporal features of functional brain networks across the lifespan. NeuroImage, 173, 498–508. 10.1016/j.neuroimage.2018.02.066

Volkow, N. D., Koob, G. F., Croyle, R. T., Bianchi, D. W., Gordon, J. A., Koroshetz, W. J., Pérez-Stable, E. J., Riley, W. T., Bloch, M. H., Conway, K., Deeds, B. G., Dowling, G. J., Grant, S., Howlett, K. D., Matochik, J. A., Morgan, G. D., Murray, M. M., Noronha, A., Spong, C. Y., … Weiss, S. R. B. (2018). The conception of the ABCD study: From substance use to a broad NIH collaboration. Developmental Cognitive Neuroscience, 32, 4–7. 10.1016/j.dcn.2017.10.002

Yarkoni, T. (2009). Big Correlations in Little Studies: Inflated fMRI Correlations Reflect Low Statistical Power—Commentary on Vul et al. (2009). Perspectives on Psychological Science, 4(3), 294–298. 10.1111/j.1745-6924.2009.01127.x

