## Supplementary Table 1 for "Metrics for estimating individual-level changes in functional brain connectivity and their correspondence with topology changes"

| ***Supplemental Table 1. Means and standard deviations for each connectivity metric, across networks.*** | | | | | |
| --- | --- | --- | --- | --- | --- |
|  | *Δr* | APΔ | \|APΔ\| | RCT | \|RCT\| |
| Aud | 0.0001 ± 0.064 | -142.8209 ± 16999.134 | 0.6074 ± 51.652 | 0.0008 ± 0.506 | 0.0104 ± 0.507 |
| between | -0.0 ± 0.063 | -159.7955 ± 17748.889 | 0.64 ± 51.665 | 0.0001 ± 0.506 | 0.0108 ± 0.507 |
| within | 0.0013 ± 0.076 | 43.8994 ± 1537.814 | 0.2489 ± 51.508 | 0.0091 ± 0.506 | 0.0063 ± 0.505 |
| COpp | 0.0001 ± 0.061 | -19.6495 ± 32121.76 | 0.5498 ± 51.317 | 0.0005 ± 0.505 | 0.007 ± 0.506 |
| between | 0.0 ± 0.061 | 112.8362 ± 23045.866 | 0.5893 ± 51.413 | 0.0 ± 0.505 | 0.0074 ± 0.506 |
| within | 0.0008 ± 0.062 | -1476.9917 ± 80865.391 | 0.1155 ± 50.265 | 0.006 ± 0.506 | 0.0027 ± 0.506 |
| CPa | -0.0 ± 0.091 | -52.6618 ± 8758.797 | 1.0441 ± 51.139 | 0.0001 ± 0.506 | 0.0144 ± 0.506 |
| between | 0.0001 ± 0.084 | -53.7393 ± 8912.793 | 1.1638 ± 51.521 | 0.0005 ± 0.506 | 0.0164 ± 0.506 |
| within | -0.0009 ± 0.148 | -40.8091 ± 6840.938 | -0.2722 ± 46.718 | -0.0036 ± 0.507 | -0.0074 ± 0.507 |
| DAttn | 0.0 ± 0.057 | -50.8745 ± 6751.68 | 0.8253 ± 51.18 | -0.0004 ± 0.506 | 0.0125 ± 0.506 |
| between | -0.0 ± 0.057 | -54.3132 ± 6969.582 | 0.935 ± 51.386 | -0.0008 ± 0.506 | 0.0142 ± 0.506 |
| within | 0.0004 ± 0.058 | -13.0481 ± 3563.73 | -0.3814 ± 48.848 | 0.0033 ± 0.506 | -0.0062 ± 0.505 |
| DMN | 0.0 ± 0.055 | 72.4537 ± 11213.242 | 1.0697 ± 51.204 | 0.0002 ± 0.506 | 0.0158 ± 0.505 |
| between | 0.0 ± 0.055 | 72.2319 ± 11696.049 | 1.0051 ± 51.253 | 0.0003 ± 0.506 | 0.0154 ± 0.505 |
| within | 0.0 ± 0.052 | 74.8931 ± 2018.315 | 1.7803 ± 50.666 | -0.0007 ± 0.506 | 0.021 ± 0.506 |
| FPN | 0.0001 ± 0.052 | 32.2784 ± 20400.583 | 1.3415 ± 51.136 | 0.0011 ± 0.506 | 0.0166 ± 0.506 |
| between | 0.0001 ± 0.052 | 34.8651 ± 21300.083 | 1.3022 ± 51.233 | 0.0004 ± 0.506 | 0.0167 ± 0.506 |
| within | 0.0009 ± 0.051 | 3.8248 ± 1894.575 | 1.7738 ± 50.063 | 0.0088 ± 0.506 | 0.0158 ± 0.505 |
| RsTp | -0.0002 ± 0.081 | 43.262 ± 6714.183 | -0.4177 ± 51.594 | -0.0018 ± 0.505 | -0.003 ± 0.505 |
| between | -0.0003 ± 0.077 | 44.2264 ± 6999.566 | -0.3967 ± 51.708 | -0.0025 ± 0.505 | -0.0022 ± 0.505 |
| within | 0.0013 ± 0.115 | 32.6538 ± 1425.425 | -0.6483 ± 50.331 | 0.0062 ± 0.504 | -0.0118 ± 0.505 |
| SMH | 0.0001 ± 0.065 | -0.6824 ± 6747.944 | 2.6441 ± 51.815 | 0.0001 ± 0.507 | 0.0336 ± 0.507 |
| between | 0.0001 ± 0.064 | 0.9526 ± 7025.228 | 2.2735 ± 51.951 | -0.0002 ± 0.507 | 0.0298 ± 0.507 |
| within | 0.0004 ± 0.081 | -18.6671 ± 1878.019 | 6.7208 ± 50.119 | 0.0032 ± 0.508 | 0.0756 ± 0.506 |
| SMM | 0.0001 ± 0.085 | 57.0387 ± 15955.651 | 0.9105 ± 51.899 | 0.0008 ± 0.506 | 0.0109 ± 0.507 |
| between | 0.0002 ± 0.078 | 62.2491 ± 16642.527 | 0.957 ± 52.076 | 0.001 ± 0.506 | 0.0122 ± 0.507 |
| within | -0.0005 ± 0.136 | -0.2757 ± 2879.202 | 0.3999 ± 49.909 | -0.0016 ± 0.507 | -0.0034 ± 0.507 |
| SN | 0.0001 ± 0.078 | -25.9344 ± 3559.804 | -0.3795 ± 51.824 | 0.0001 ± 0.507 | -0.0028 ± 0.506 |
| between | -0.0 ± 0.074 | -31.6095 ± 3697.891 | -0.3855 ± 51.821 | -0.0003 ± 0.507 | -0.0023 ± 0.506 |
| within | 0.0011 ± 0.118 | 36.4921 ± 1282.477 | -0.3134 ± 51.867 | 0.0043 ± 0.506 | -0.0088 ± 0.507 |
| VAttn | 0.0 ± 0.055 | 93.6414 ± 22287.186 | 0.0614 ± 51.152 | 0.0003 ± 0.506 | 0.0013 ± 0.506 |
| between | 0.0 ± 0.055 | 109.1301 ± 23244.015 | 0.0519 ± 51.29 | 0.0002 ± 0.506 | 0.0011 ± 0.506 |
| within | 0.0003 ± 0.054 | -76.7342 ± 4181.668 | 0.1649 ± 49.624 | 0.0015 ± 0.505 | 0.0035 ± 0.505 |
| Vis | -0.0002 ± 0.07 | -93.1177 ± 16025.237 | 0.3074 ± 51.569 | -0.0011 ± 0.506 | 0.0088 ± 0.506 |
| between | -0.0002 ± 0.067 | -102.6152 ± 16725.717 | 0.3188 ± 51.658 | -0.001 ± 0.506 | 0.0091 ± 0.506 |
| within | -0.0004 ± 0.095 | 11.3546 ± 2109.633 | 0.1818 ± 50.587 | -0.0027 ± 0.506 | 0.006 ± 0.505 |
| *Note*. (A) Twelve functional networks were defined using resting-state functional connectivity patterns according to methods described by Gordon et al. (2016). (B) Networks’ within connectivity measures a network’s connection strength with that network. Networks’ between connectivity measures a network’s connection strength with other networks, (C) Functional connectivity change metrics were chosen based on methods described by Bottenhorn et al., (2025). | | | | | |
